## Supporting information for "Transcriptome-wide identification of 5-methylcytosine by deaminase and reader protein-assisted sequencing"

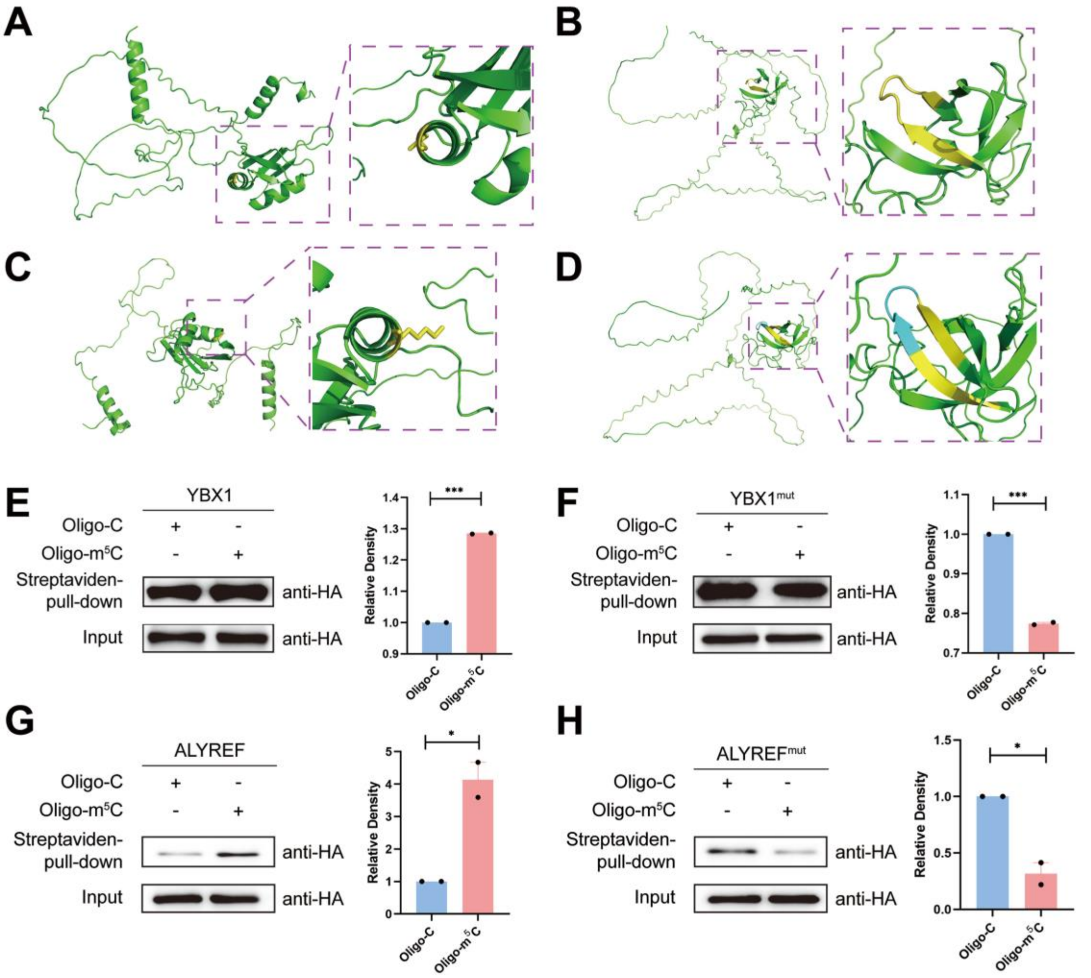


**Supplementary Figure.1: The 3D structures of ALYREF$^{K171A}$, YBX1$^{KO\ W65\text{-}N70}$, ALYREF and YBX1.**

**(A, B)** Computer simulation of the three-dimensional structure of mutated ALYREF (**A**) and YBX1 (**B**) protein. On the left is the full structural model of ALYREF$^{K171A}$, and on the right the 171st amino acid is colored in yellow and shows the stick structure of mutated A171 (**A**). The complete structural model of YBX1 after knockdown of W65-N70 (WFNVRN) is shown on the left, and the original L60-K64 and G71-I75 amino acids are colored in yellow on the right (**B**).

**(C, D)** Computer simulation of the three-dimensional structure of wild-type ALYREF(**C**) and YBX1(**D**). The stick structure of K171 is highlighted in yellow(**C**). The amino acids L60-K64 and G71-I75 is marked in yellow, while the W65-N70 (WFNVRN) region is highlighted in blue**(D)**.

**(E-H)** The RNA pulldown assay demonstrates the binding of expressed YBX1(**E**), YBX1$^{mut}$(**F**), ALYREF(**G**) and ALYREF$^{mut}$(**H**) proteins to biotinylated RNA containing C or m$^5$C. The corresponding quantitative analyses of their binding capacities are shown on the right.

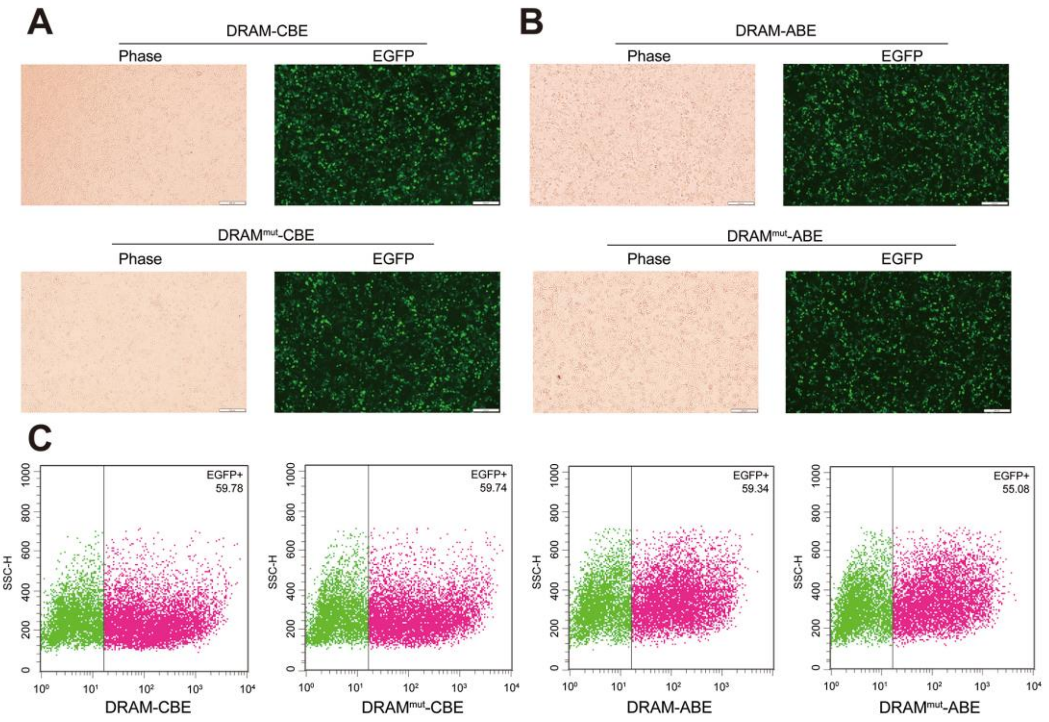


**Supplementary Figure.2: Expression of DRAM and DRAM$^{mut}$ in HEK293T cells.**
**(A, B)** Examination of the EGFP-positive cells by fluorescence microscopy. Scale bars, 200μm.
**(C)** Quantification of the EGFP-positive cells by flow cytometry.

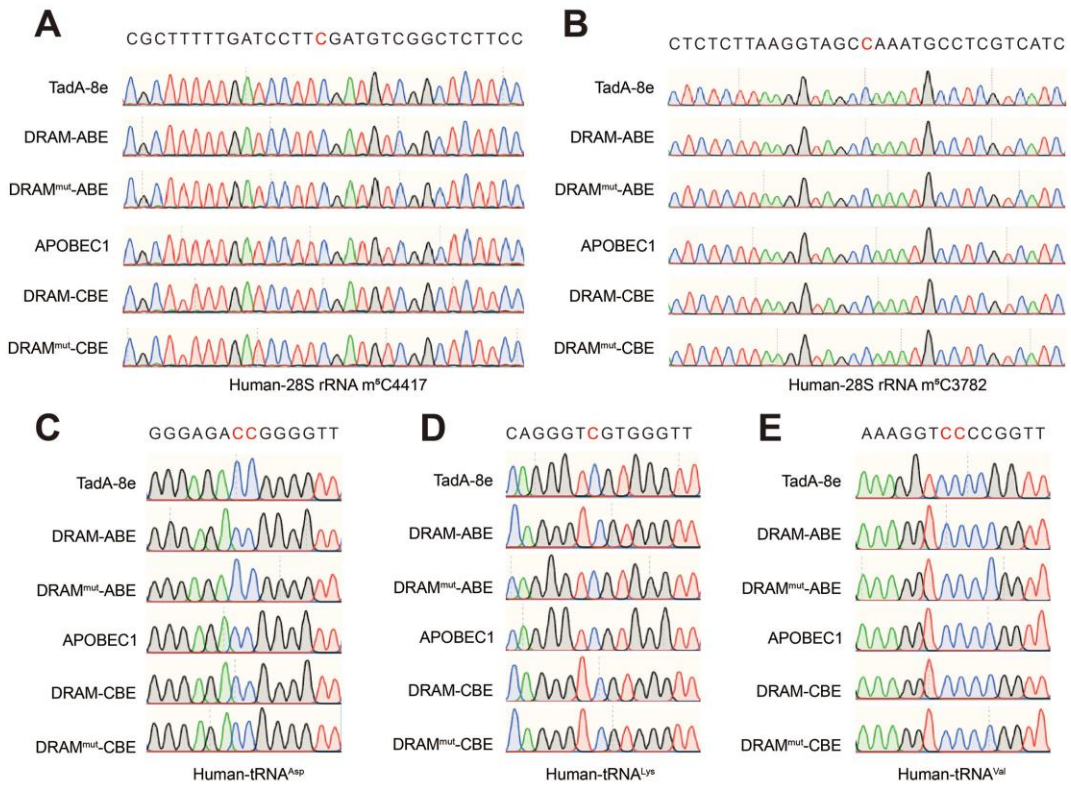


**Supplementary Figure.3: Editing of the DRAM system near the $m^5C$ sites on 28S rRNA and tRNA.**

RT-PCR followed by Sanger sequencing verified the presence of $m^5C$ modifications at specific sites in DRAM-transfected HEK293T cells, including $m^5C4447$ in 28S rRNA (**A**), $m^5C3782$ in 28S rRNA (**B**), $m^5C48$ and $m^5C49$ in $tRNA^{Asp}$ (**C**), $m^5C48$ in $tRNA^{Lys}$ (D), and $m^5C48$ and 49 in $tRNA^{Val}$ (**E**). Mutations were observed in sequences flanking the $m^5C$ loci.

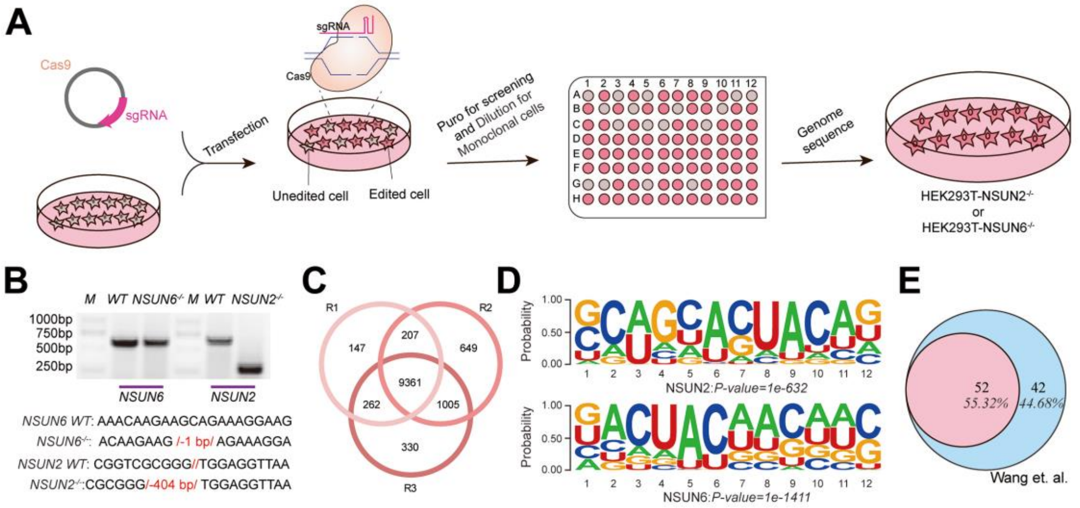


**Supplementary Figure.4: CRISPR/Cas9 -mediated depletion of NSUN2 and NSUN6 in HEK293T.**

**(A)** Schematic representation the construction of NSUN2$^{-/-}$ and NSUN6$^{-/-}$ Cell line. HEK293T cells were transfected with plasmids encoding a PX459 editor and guide RNA targeting the NSUN2 or NSUN6 gene. After puro for screening, cells were diluted and seeded into a 96-well plate to allow for clonal selection. Monoclonal cells were then picked and screened for the desired genotype, resulting in the identification of the NSUN2$^{-/-}$ and NSUN6$^{-/-}$ Cell line.

**(B)** Mutation detection of NSUN2 and NSUN6 by PCR. M, the marker used is DL2000.

**(C**) Venn diagram showing the overlap of DRAM-edited mRNAs identified across three sets of biological replicates.

**(D)** Motif enrichment analysis based on the sequences spanning 10 nt upstream and downstream of DRAM-edited sites mediated by loci associated with NSUN2 or NSUN6.

**(E)** Stacked Venn diagram showing the overlap of genes detected by *Wang et al.* and those identified by the DRAM system.

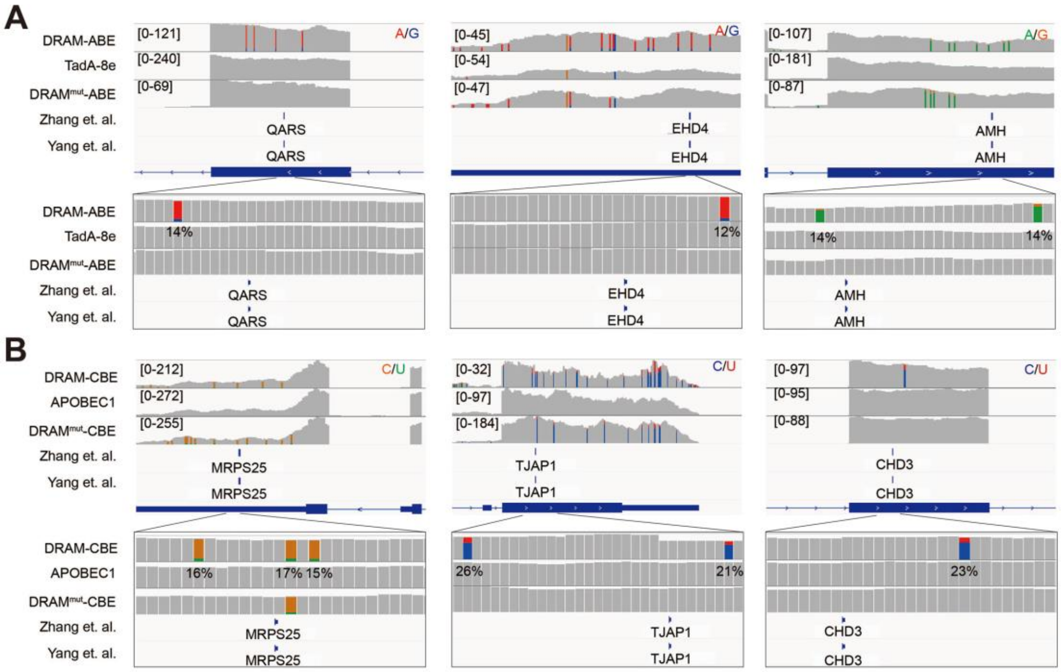


**Supplementary Figure.5: DRAM-Seq editing in cellular RNAs.**

Integrative genomics viewer (IGV) browser traces of DRAM-seq data expressing the indicated constructs in QARS(**A**), EHD4(**A**), AMH (**A**), MRPS25 (**B**), TJAP1 (**B**) and CHD3 (**B**) mRNAs. C-to-U or A-to-G mutations found in at least 10% of reads are indicated by coloring. The previously published RNA BS-seq datasets from two individual studies were displayed as panel "Yang et al." and "Zhang et al.".

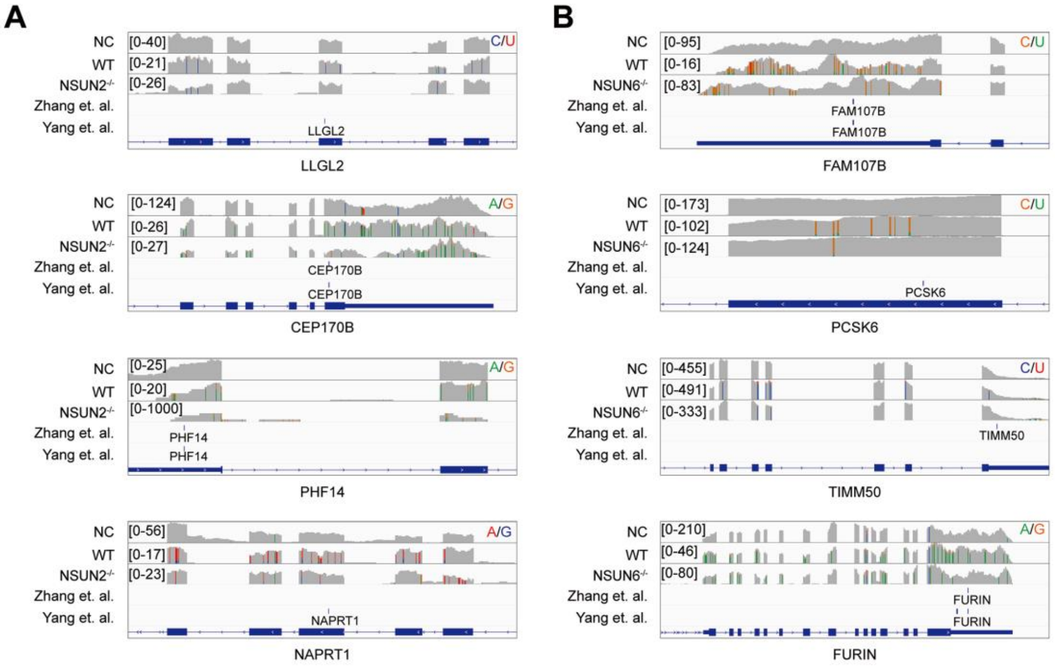


**Supplementary Figure.6: DRAM-Seq analysis in NSUN2-depleted and NSUN6-depleted cellular RNAs.**

IGV Browser traces showing read coverage and C-to-U/A-to-G mutations for eight representative mRNAs in wild-type, NSUN2-depleted cells (**A**) and NSUN6-depleted cells (**B**): LLGL2, CEP170B, PHF14, NAPRT1, FAM107B, PCSK6, TIMM50 and FURIN. C-to-U or A-to-G mutations were found in at least 10% of reads are indicated by coloring. The previously published RNA BS-seq datasets from two individual studies were displayed as panel "Yang et al." and "Zhang et al.".

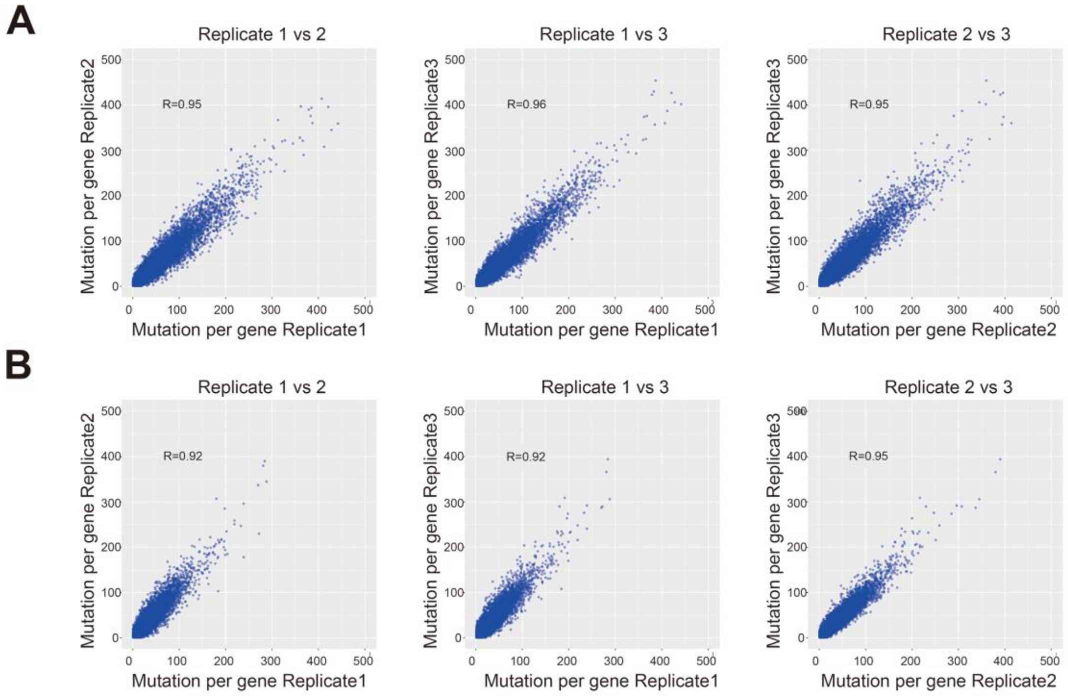


**Supplementary Figure.7: Analysis of DRAM-seq data repeatability.**

Three individual DRAM-ABE replicates were analyzed for the number of A-to-G mutations per RNA(**A**). Three individual DRAM-CBE replicates were analyzed for the number of C-to-U mutations per RNA(**B**). Pearson correlation coefficient indicated a high degree of overlap between individual replicates.

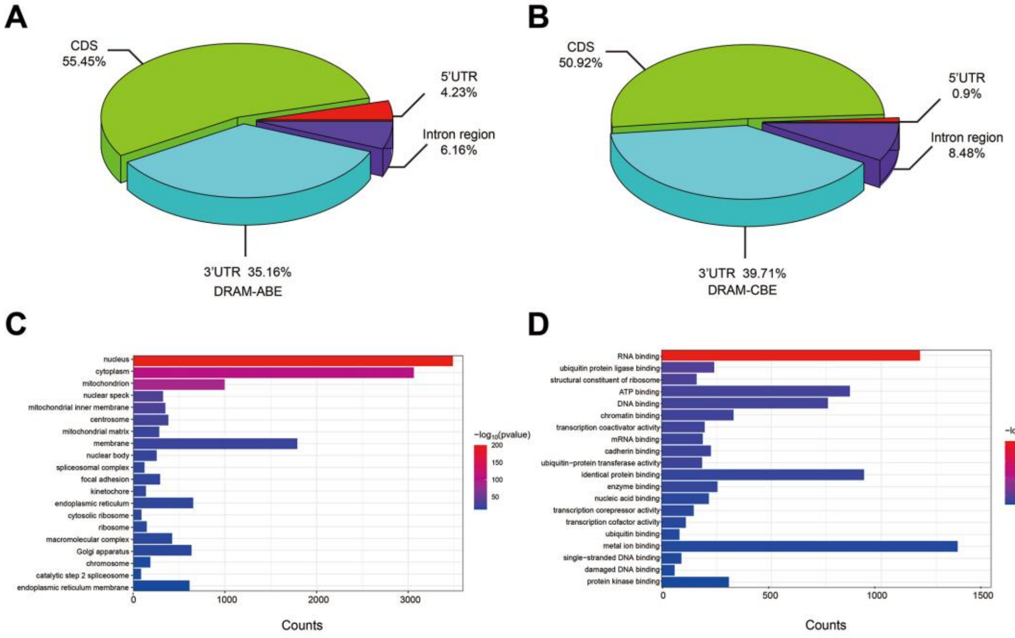


**Supplementary Figure. 8：Transcriptome-Wide mapping of m$^5$C in HEK293T cells.**

(**A**, **B**) The pie chart shows the distribution of editing sites in different transcript regions in cells expressing DRAM-ABE and DRAM-CBE.

(**C**, **D**) Gene Ontology (GO) cellular component(**C**) and molecular function(**D**) enrichment analysis of genes with DRAM-Seq editing events. Statistical analyses were performed using the DAVID tool. $p < 0.05$.

**Supplementary Table 1: The detection for $m^5C$ in RNA**

| Detection method | Advantages | Disadvantages |
|---|---|---|
| BS-seq | Single nucleotide resolution; Primitive methylation patterns. | Influenced by the secondary structure of RNA; Disability for distinguishing $m^5C$ from other type of cytosine modifications, Causes RNA damage. |
| $m^5C$-RIP-seq | Distinguishable between $m^5C$ and other types of cytosine modifications. | Insensitive to low-abundance $m^5C$; Influenced by the secondany structure of RNA; Relatively low esolution; Highly dependent on antibody specificity. |
| miCLIP-seq | Specific detection of the $m^5C$ sites of NSUN2 methyltransferase. | Unable to detect $m^5C$ sites of other methyltransferases; Highly dependent on antibody specifity; Altered methylation patterns; Time-consuming; Expensive. |
| AZA-lP-seq | Methyltransferase-specific detection of $m^5C$; Single nucleotide resolution. | Highly dependent on antibody specificity and 5-azaC binding efficiency; High toxiciticy of 5-azaC. |
| TAWO-seq | Single nucleotide resolution; Non-methylated cytosines have a low false positive rate of conversion. | Unstable conversion efficiency; Dependent on the oxidation efficiency of peroxotungstate. |
| Nanopore-seq | Raw DNA or RNA can be directly sequenced. | Expensive; High error rate; Sequencing signals probably affected by multiple nucleotides at the same time. |

**Supplementary Table 4:**

**The primers for vector construction**

| Name of Primer | Sequences (5′−3′) |
|---|---|
| ABE-ALYREF-ALYREF-Fwd | agcagcggggggtcaatgcccgattccgcg |
| ABE-ALYREF-ALYREF-Rev | ttttgagccgccagaactggtgtccattctcgcat |
| ABE-ALYREF-TadA-8e-Fwd | tctggcggctcaaaaagaacc |
| ABE-ALYREF-TadA-8e-Rev | tgaccccccgctgctg |
| ABE-YBX1-YBX1-Fwd | agcagcggggggtcaatgagcagcgaggccgagacccagc |
| ABE-YBX1-YBX1-Rev | ttttgagccgccagactcagccccgccctgc |
| ABE-YBX1-TadA-8e-Fwd | tctggcggctcaaaaagaacc |
| ABE-YBX1-TadA-8e-Rev | tgaccccccgctgctg |
| CBE-ALYREF-ALYREF-Fwd | gagtccgccacaccaatgcccgattccgcg |
| CBE-ALYREF-ALYREF-Rev | gtaggggtactcgagactggtgtccattctcgcattatagg |
| CBE-ALYREF-APOBEC1-Fwd | ctcgagtacccctacgacgtg |
| CBE-ALYREF-APOBEC1-Rev | tggtgtggcggactctg |
| CBE-YBX1-YBX1-Fwd | gagtccgccacaccaatgagcagcgaggccgagac |
| CBE-YBX1-YBX1-Rev | gtaggggtactcgagctcagccccgccctg |
| CBE-YBX1-APOBEC1-Fwd | ctcgagtacccctacgacgtg |
| CBE-YBX1-APOBEC1-Rev | tggtgtggcggactctg |
| YBX1$^{mut}$-Fwd | ttgtacaccagactgccataaggaagtaccttcgcagtgt |
| YBX1$^{mut}$ -Rev | acactgcgaaggtacttccttatggcagtctggtgtacaa |
| ALYREF $^{mut}$-Fwd | gccctgaaggccatggcgcagtacaacggcgt |
| ALYREF $^{mut}$ -Rev | acgccgttgtactgcgccatggccttcagggc |

**The primers used for genotyping**

| Name of Primer | Sequences (5′−3′) |
|---|---|
| NSUN2-KO -PCR-Fwd | ccccttagagctgttcgctgt |
| NSUN2-KO -PCR-Rev | gtcgaagaaaacaccgtcgcctt |
| NSUN6-KO -PCR-Fwd | taccatgttgaagcccaagaa |
| NSUN6-KO -PCR-Rev | gaagctactaaggcccagttt |

**Single guide RNA**

| Name of Primer | Sequences (5′−3′) |
|---|---|
| NSUN2-PX459sgRNA1-H-Fwd | caccgcgccatcctccgcgtcctc |
| NSUN2-PX459sgRNA1-H-Rev | aaacgaggacgcggaggatggcgc |
| NSUN2-PX459sgRNA2-H-Fwd | caccgggtggtggaaagcgcggcg |
| NSUN2-PX459sgRNA2-H-Rev | aaaccgccgcgctttccaccaccc |
| NSUN2-PX459sgRNA3-H-Fwd | caccgaggctaccccgagatcgtca |
| NSUN2-PX459sgRNA3-H-Rev | aaactgacgatctcggggtagcctc |
| NSUN2-PX459sgRNA4-H-Fwd | caccgtgttctccttgacgatctcg |
| NSUN2-PX459sgRNA4-H-Rev | aaaccgagatcgtcaaggagaacac |

| | |
|---|---|
| NSUN6-PX459sgRNA1-H-Fwd | cacctaggtaaacaagaagcagaa |
| NSUN6-PX459sgRNA1-H-Rev | aaacttctgcttcttgtttaccta |
| NSUN6-PX459sgRNA2-H-Fwd | caccatttttcacatgttgtactg |
| NSUN6-PX459sgRNA2-H-Rev | aaaccagtacaacatgtgaaaaat |

**The primers for DRAM-Sanger analysis**

| Name of Primer | Sequences (5′−3′) |
|---|---|
| RPSA-DRAM-Few | gcaaatgaaggaggaggatgt |
| RPSA-DRAM-Rev | gttagtgaaggttccaggagtg |
| AP5Z1-DRAM-Few | agggacttcggtgcagatta |
| AP5Z1-DRAM-Rev | ctcaagcctccaatcagagc |
| tRNA-Val-DART- Few | gtttccgtagtgtagtgg |
| tRNA-Val-DART-Rev | ctcaactggtgtcgtggagtcggcaattcagttgagtggtgtttccgccc |
| tRNA-Lys-DART- Few | gcccggctagctcagt |
| tRNA-Lys-DART-Rev | ctcaactggtgtcgtggagtcggcaattcagttgagtggcgcccaacgtg |
| tRNA-Asp-DART- Few | tcctcgttagtatagtg |
| tRNA-Asp-DART-Rev | ctcaactggtgtcgtggagtcggcaattcagttgagtggtggctccccgt |
| 28S rRNA4417-DART- Few | ggtacacctgtcaaacggtaa |
| 28S rRNA4417-DART-Rev | ccaagcacatacaccaaatgtc |
| 28S rRNA3782-DART- Few | cagccgacttagaactg |
| 28S rRNA3782-DART-Rev | cctcccacttattctacac |

**The primers for qPCR**

| Name of Primer | Sequences (5′−3′) |
|---|---|
| NSUN2-H-qPCR-Fwd | atcttgagaaaatcgccacac |
| NSUN2-H-qPCR-Rev | atcattcgcaataacaaatccct |
| NSUN6-H-qPCR-Fwd | tcagcgtgatcggcaagatt |
| NSUN6-H-qPCR-Rev | acctaaagcagtcacaatctcct |
| GAPDH-H-qPCR-Fwd | gtctcctctgacttcaacagcg |
| GAPDH-H-qPCR-Rev | accaccctgttgctgtagccaa |

**The primers for bisulfite sequencing PCR**

| Name of Primer | Sequences (5′−3′) |
|---|---|
| RPSA-BS-Fwd | tgttgttattgaaaattttgttgatg |
| RPSA-BS-Rev | ataatcaaccctaaaatcaataacca |
| AP5Z1-BS-Fwd | gttagtttgttattgaggttaggtt |
| AP5Z1-BS-Rev | ctcaactcaatattacctaaaaacaaa |
